# Interleaved intersectional strategy enables genetic lineage tracing with enhanced specificity

**DOI:** 10.1101/2024.03.06.583635

**Authors:** Maoying Han, Zhicong Liu, Xiuzhen Huang, Lei Liu, Bo Zhou, Kathy O. Lui, Qiang Shu, Bin Zhou

## Abstract

**BACKGROUND:** Dual recombinases have been increasingly employed for enhanced precision in genetic targeting. A recent study utilizing an intersectional genetic approach through dual recombinases (Dre + CreER) has revealed that endocardium-derived fibroblasts (EndoFbs) play a pivotal role in cardiac fibrosis after pressure overload. However, this intersectional strategy has limitations, primarily due to ectopic genetic labeling of non-target cells within the adult heart by the constitutively active Dre recombinase.

**METHODS:** To address this issue, we have developed an advanced, interleaved and intersectional reporter (IIR) strategy in this study. This IIR strategy leverages an inducible CreER to prevent inadvertent Dre-rox recombination during development or disease progression by designing an interleaved reporter to allow for more specific tracing of EndoFbs. Moreover, our IIR system also incorporates Diphtheria Toxin Receptor (DTR) in targeted cells, enabling functional characterization of these cells after genetic ablation.

**RESULTS:** EndoFbs were regionally distributed in the heart during homeostasis and proliferated preferentially in response to pressure overload, leading to cardiac fibrosis in defined regions. The IIR strategy enables the tracing of EndoFbs with a more prominent regional pattern and facilitates genetic ablation of EndoFbs through DT injection. In addition, we have applied this IIR strategy to specifically target fibroblasts derived from the epicardium (EpiFbs). Genetic lineage tracing of EpiFb reveals that their distribution pattern is complementary to that of EndoFbs in the adult heart. When a substantial number of EpiFbs were genetically ablated, EndoFbs could replace the loss of EpiFbs in some specific regions of hearts.

**CONCLUSIONS:** The IIR strategy refines the precision of genetic lineage tracing while still employing the constitutively active Dre recombinase in tandem with inducible Cre. EndoFbs and EpiFbs are complementary in their distribution pattern in the heart, where EndoFbs have the potential to replace the loss of EpiFbs in some regions.

## INTRODUCTION

Cre-loxP system has been widely used for cell fate mapping and genetic manipulations in multiple biomedical research.^1–3^ The precision of cell lineage tracing studies largely depends on the fidelity of the gene promoter that drives Cre recombinase, which is not always exclusively specific to the target cell type as expected.^4^ The use of non-specific lineage tracing tools has led to inconclusive findings over the cellular origin and fate plasticity in multiple biomodel research fields.^5–9^ For example, earlier works have acknowledged c-Kit as a stem cell marker of cardiomyocytes;^5, 10^ however, recent lineage tracing data showed that it is also expressed by cardiomyocytes,^11^ questioning the existence of cardiac stem cells proposed for decades. Based on the orthogonal features of different recombinases, lineage tracing with dual recombinases driven by two different gene promoters enables cell labeling with two markers and, therefore, enhances the precision of targeting a specific cell lineage.^12–14^ Dual recombinases-mediated lineage tracing technology undoubtedly demonstrates the lack of cardiac stem cells in adult mammalian heart.^15^ Similarly, whether PDGFRb^+^ pericytes or PDGFRa^+^ fibroblasts behave as mesenchymal stem cells *in vivo* remain controversial.^16, 17^ The generation of *Pdgfra-DreER* and *Pdgfrb-CreER* mouse lines enables tracing of these cells in the same mouse through intersectional genetics.^18^ The fate mapping study has revealed that PDGFRa^+^PDGFRb^+^ cells preferentially contribute to *de novo* adipogenesis in adult mice, redefining the origin of intramyocardial adipocytes.^18, 19^ The dual recombinases-mediated lineage tracing strategy begins to be broadly applied to multiple research fields for enhancing the specificity of cell target or for targeting multiple cell lineages in the same *in vivo* system.^20–27^

More recently, dual recombinases are also used to specifically demonstrate the contribution of a fibroblast subset to cardiac fibrosis in the adult myocardium.^28^ Excessive cardiac myofibroblasts in disease are derived from pre-existing fibroblasts through expansion, which are key mediators of tissue fibrosis and determinants of scar formation.^29, 30^ Cardiac fibroblasts residing in different parts of the myocardium are heterogeneous as the diverse developmental origins from the epicardium, endocardium, and neural crest all contribute to cardiac fibroblasts.^9, 31–34^ Specifically, endocardial cells undergo endothelial-to-mesenchymal transition to give rise to a subset of cardiac fibroblasts located in distinct regions of the heart during development.^9, 35^ Since endocardial cells also generate coronary vascular endothelial cells, pericytes, smooth muscle cells, and adipocytes, in addition to endocardium-derived fibroblasts (EndoFbs),^36–39^ EndoFbs cannot be specifically targeted using a single endocardial cell marker. Specific tracing of these cells is achieved with both endocardial cell marker Nfatc1 and fibroblast marker Col1a2 driven Dre and CreER, respectively.^28^ Using intersectional reporter *Rosa26-rox-Stop-rox-loxP-Stop-loxP-GFP* (*R26-RL-GFP*), Dre-rox recombination in endocardial cells of *Nfatc1-Dre*; *Col1a2- CreER*; *R26-RL-GFP* mice removes the first Stop cassette flanked by two rox sites, and subsequent tamoxifen (Tam)-induced *Col1a2-CreER* excises the second Stop cassette flanked by two loxP sites, leading to GFP expression only in EndoFbs in adult hearts (Figure 1A). This strategy is useful for genetically distinguishing a particular cell subset of one origin from other subsets derived from multiple developmental origins within a heterogeneous cell population.

**Figure 1.**
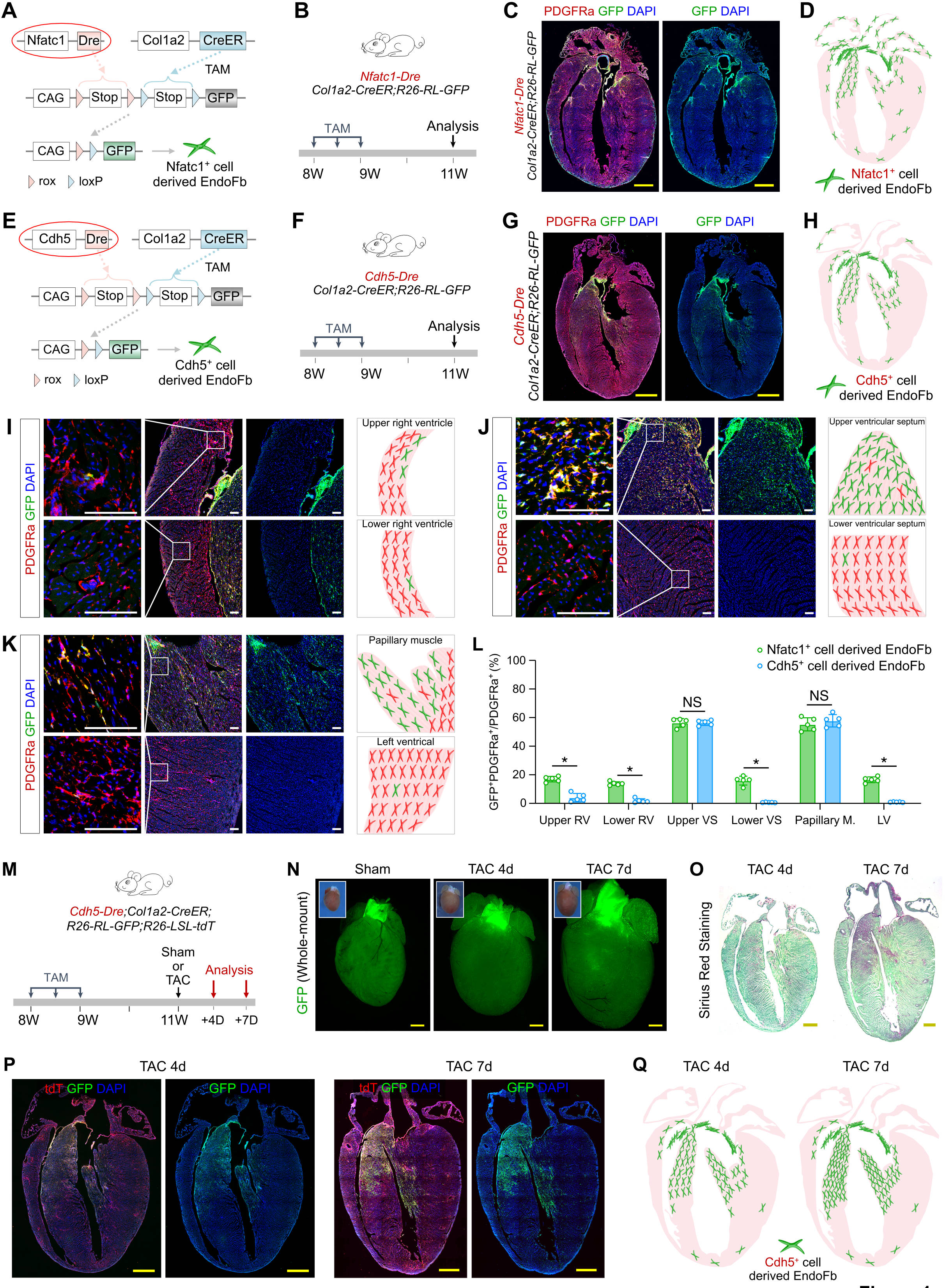
Tracing EndoFbs by *Nfatc1-Dre* and *Cdh5-Dre*. A, A schematic showing the strategy for genetic tracing of EndoFbs by *Nfatc1-Dre;Col1a2-CreER;R26-RL-GFP* mice. **B,** A schematic showing the experimental design. **C,** Immunostaining for PDGFRα and GFP on adult heart sections. **D,** A cartoon image showing the distribution of Nfatc1-derived EndoFbs in the adult heart. **E**, A schematic showing the strategy for genetic tracing of EndoFbs by *Cdh5- Dre;Col1a2-CreER;R26-RL-GFP* mice. **F,** A schematic showing the experimental design. **G,** Immunostaining for PDGFRα and GFP on adult heart sections. **H,** A cartoon image showing the distribution of Cdh5-derived EndoFbs in the adult heart. **I-K,** Immunostaining for PDGFRα and GFP on different parts of adult heart sections collected from *Cdh5-Dre;Col1a2-CreER;R26-RL- GFP* mice. The boxed regions are magnified in the panels on the left, and the right cartoon shows the distribution of EndoFbs in the indicated regions of hearts. **L,** Quantification of the percentage of PDGFRα^+^ EndoFbs expressing GFP in different regions of the heart. **M**, A schematic showing the experimental design. **N,** Whole-mount fluorescence views of hearts collected from *Cdh5-Dre;Col1a2-CreER;R26-RL-GFP;R26-LSL-tdT* mice under sham treatment or at 4 days or 7 days after TAC. The inserts indicate bright-field images. **O,** Sirius Red staining of heart sections. **P,** Immunostaining for tdT and GFP on heart sections from mice at 4 days or 7 days after TAC. **Q,** A cartoon image showing the distribution of Cdh5-derived EndoFbs in hearts at 4 days or 7 days after TAC. RV, right ventricle; LV, left ventricle; VS, ventricular septum; Papillary M., papillary muscle. The quantification data is represented as the mean ± SD; \**P* < 0.05; NS, non-significant; n = 5 mice. Scale bars: yellow, 1 mm; white, 100 µm.

However, the combinatorial use of dual recombinases such as Dre and CreER for genetic lineage tracing still exhibits some caveats. For instance, if Nfatc1 is transiently expressed in non- EndoFbs during development, the unexpected despite transient activation of *Nfatc1-Dre* would result in Dre-rox recombination and, therefore, ectopic labeling of other cardiac fibroblasts in addition to EndoFbs. Such ectopic labeling is often technically hard to discern labeled cells as solely EndoFbs because it requires a positive control to confirm the specific labeling pattern of EndoFbs ideally with an endocardium lineage-restricted Dre driver that is not activated in other cell populations of the adult heart at any time. The current study aims to provide an advanced strategy to resolve the ectopic labeling issue associated with the use of a constitutively active recombinase in dual recombinases-mediated genetic lineage targeting studies. We iterated the intersectional genetic approach (loxP-Stop-loxP-rox-Stop-rox-reporter) by integrating the strategy of using an interleaved reporter (loxP-rox-loxP-rox-reporter), and developed an interleaved intersectional reporter (IIR) system for specific labeling of cell populations by using Dre & CreER recombinases.

## METHODS

### Mice

All mice mentioned in this study were used in accordance with the guidelines set forth by the Institutional Animal Care and Use Committee (IACUC) at the Institute of Biochemistry and Cell Biology, Center for Excellence in Molecular Cell Science, Chinese Academy of Science. The *Nfatc1-Dre*,^37^ *Col1a2-CreER*,^40^ *R26-RL-GFP*,^28^ *Cdh5-Dre*,^41^ *R26-LSL-tdTomato*,^42^ and *Upk3b- Dre*^18^ mouse lines were reported previously. The *IIR* mouse line (Shanghai Biomodel Organism) was generated by homologous recombination using CRISPR-Cas9 technology. In brief, a cDNA encoding CAG-loxP-Stop-loxP-rox-loxP-GFP-polyA-rox-tdTomato-P2A-DTR-WPRE-ployA was inserted between the exons 1 and 2 of the endogenous Rosa26 gene locus to generate the *IIR* mouse line. All mice were maintained under a mixed C57BL6/ICR background and subjected to a 12-hour light/12-hour dark cycle. They were fed on standard chow diet (Jiangsu Xietong Pharmaceutical Bioengineering, 1010085), and were randomly assigned to experimental groups.

### Tamoxifen Treatment

Tamoxifen (Tam, Sigma, T5648) was dissolved in corn oil at 20 mg/mL as stock. A dose of 0.2 mg/g of body weight was administered via oral gavage at designated time points to activate CreER activity. For *Nfatc1-Dre;Col1a2-CreER;R26-RL-GFP* and *Cdh5-Dre;Co11a2- CreER;R26-RL-GFP* mice, three doses of Tam were consecutively administered at 8 weeks of age to enhance CreER-mediated recombination efficiency. For *Nfatc1-Dre;Col1a2-CreER;IIR* and *Upk3b-Dre;Col1a2-CreER;IIR* mice, a single dose of Tam was administered at postnatal day 14.

### Diphtheria Toxin Treatment

Diphtheria toxin (DT, Sigma, D0564-1MG) was dissolved in PBS at 1 µg/µl as stock. A dose of 10 µg/g of body weight was administered via intraperitoneal injection at the assigned time points to genetically ablate target cells. For *Nfatc1-Dre;Col1a2-CreER;IIR* mice, three doses of DT were administered to ablate EndoFbs. Similarly, three doses of DT were administered to *Upk3b- Dre;Col1a2-CreER;IIR* mice to ablate EpiFbs.

### Genomic Polymerase Chain Reaction

Genomic DNA was extracted from the mouse toes using a lysis buffer consisting of 5 mmol/L EDTA, 100 mmol/L Tris HCl at pH 7.8, 200 mmol/L NaCl, 0.2% sodium dodecyl sulfate, and 100 µg/mL proteinase K at 55 °C overnight. The lysate was thoroughly mixed using vortex agitation to ensure complete dissolution in the lysis buffer. DNA was precipitated by adding an equal volume of absolute ethanol, followed by centrifugation at 15,000 rpm for 3 minutes. The resulting pellet was washed with 70% ethanol, and the DNA was subsequently dissolved in distilled H2O. Genotyping of all mice was performed using genomic polymerase chain reaction (PCR) with specific primers designed using MacVector.

### Tissue Collection and Whole-mount Fluorescence Microscopy

The target tissues were harvested from mice with specific genotypes and subsequently washed in cold phosphate-buffered saline (PBS; Gibco, C10010500BT) to remove blood. Next, the tissues were fixed with 4% paraformaldehyde at 4 °C for 1 hour. After fixation, the tissues were washed with cold PBS for three times. They were then oriented and placed in a Petri dish containing 1% agarose gel supplemented with PBS. Whole-mount bright-field and fluorescence images were captured using the Zeiss stereo microscope (AxioZoom V16).

### Immunostaining and Confocal Microscopy

All tissues subjected for immunostaining were initially dehydrated overnight in 30% sucrose solution. Following dehydration, the tissues were embedded in OCT (Sakura) for 1 hour at 4 °C and subsequently freezed at –20 °C. Cryosections (thickness, 10 µm) were collected from the selected tissues at the appropriate cutting temperature and adhered to negatively charged slides. The sections were then air-dried in the fume hood for 1 hour at room temperature. After drying, the sections were washed three times with PBS for 5 minutes each. Afterwards, the sections were blocked with 2.5% normal donkey serum (Jackson Immunoresearch) and Diamidino-2- phenylindole (DAPI) (1:1,000) (Vector Labs) in PBST (0.2% Triton X-100 in PBS) for at least 30 minutes at room temperature. Following blocking, the sections were incubated with primary antibodies overnight at 4 °C, followed by three washes with PBS for 5 minutes each. Alexa- conjugated secondary antibodies (Invitrogen) were then applied to the sections and incubated for 30 minutes at room temperature. Finally, the sections were washed three times with PBS and mounted using mounting medium. The antibodies used in this study are listed: primary antibodies including PDGFRa (R&D, AF1062, 1:100), GFP (Abcam, ab6662, 1:500), tdT (Rockland, 600-401-379, 1:1,000), tdT (Proteintech, 5F8, 1:100), human HB-EGF (DTR) (R & D, AF-259-NA, 1:100); and secondary antibodies including Alexa donkey anti-rabbit 555 (Invitrogen, A31572, 1:1,000), Alexa donkey anti-rabbit 488 (Invitrogen, A21206, 1:1,000), Alexa donkey anti-goat 488 (Invitrogen, A11055, 1:1,000), Alexa donkey anti-goat 555 (Invitrogen, A32816, 1:1,000), Alexa donkey anti-goat 647 (Invitrogen, A21447, 1:1,000); Alexa donkey anti-rabbit 647 (Invitrogen, A31573, 1:1,000); Alexa donkey anti-rat 594 (JIR, 712-585- 153, 1:1,000). Immunostaining pictures were obtained by Nikon confocal laser scanning microscope (Nikon A1 FLIM).

### Sirius Red Staining

Cardiac fibrosis was assessed by Sirius Red staining. Heart sections were fixed in 4% paraformaldehyde for 15 minutes, followed by three washes with PBS for 5 minutes each. Subsequently, sections were fixed for 24 hours in Bouin solution (composed of 9% formaldehyde, 5% acetic acid, and 0.9% picric acid). On the following day, sections were washed with tap water and then immersed in 0.1% Fast Green solution (Fisher) for 3 minutes. After incubation, sections were rinsed with tap water and treated with 1% acetic acid for 1 minute. Following another wash with distilled water, sections were stained with 0.1% Sirius Red solution for 2 minutes, followed by a rinse with distilled water. Subsequently, sections were dehydrated in 95% ethanol, followed by 100% ethanol, and finally xylene, with each step lasting for 5 minutes twice. Lastly, the sections were mounted with a resinous medium, and images were captured using an Olympus microscope (model BX53).

### Statistical Analysis

The data were presented as means ± SD of at least 5 biological duplicates, each performed independently under the same circumstance. For each mouse heart or liver, 8-16 sections were collected for immunostaining, and from each section, 3-5 fields were selected for quantification. The researcher who quantified the fluorescent staining images was blinded to the groups of mice. Multiple unpaired t-test was used to compare the means of two independent groups in Figure 1, panel L; Figure 3, panel H; Figure 4, panel H and I. Two-way ANOVA followed by Tukey’s multiple comparisons test was used to compare means among multiple groups in Figure 2, panel H. Unpaired student t-test was used to compare the means of two groups in Figure 3, panel F; Figure 4, panel F. *P* <0.05 was considered as statistically significant.

**Figure 2.**
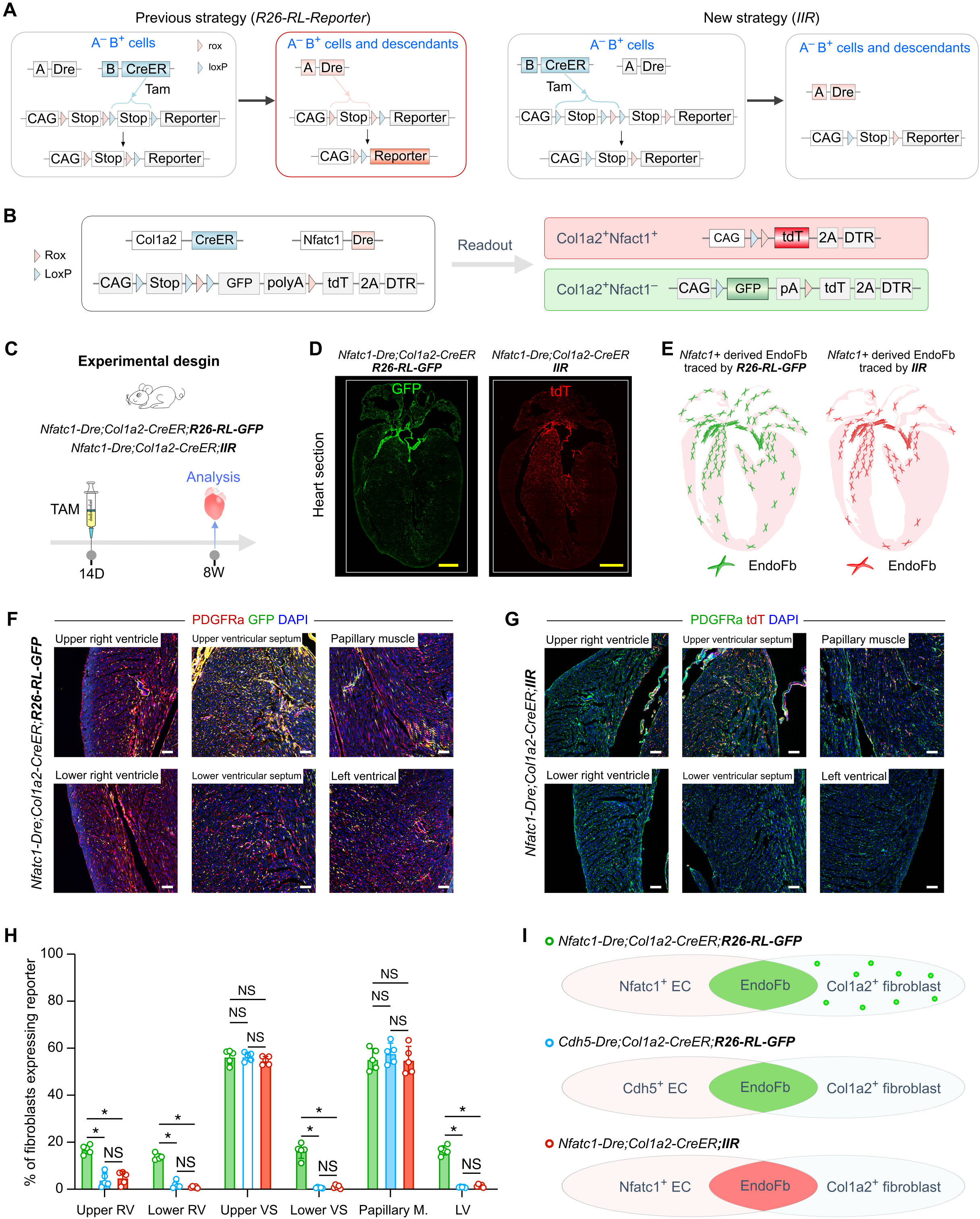
The IIR strategy effectively mitigates ectopic lineage tracing. A, Schematic diagrams showing the ectopic lineage tracing by A-Dre using *R26-RL-Reporter* (left) and prevention of Dre-rox recombination using IIR strategy (right). **B,** A schematic showing fluorescent reporter readout of EndoFbs and other fibroblast populations by using *Col1a2- CreER;Nfatc1-Dre;IIR* mice. **C,** A schematic showing the experimental design. **D,** Immunostaining for GFP or tdT on adult heart sections collected from *Nfatc1-Dre;Col1a2- CreER;R26-RL-GFP* mice (left) or *Nfatc1-Dre;Col1a2-CreER;IIR* mice (right). **E**, Cartoon images showing the distribution of EndoFbs in the adult hearts by using two different tracing strategies. **F,** Immunostaining for PDGFRα and GFP on different parts of adult heart sections collected from *Nfatc1-Dre;Col1a2-CreER;R26-RL-GFP* mice. **G**, Immunostaining for PDGFRα and tdT on different parts of adult heart sections collected from *Nfatc1-Dre;Col1a2-CreER;IIR* mice. **H**, Quantification of the percentage of PDGFRα^+^ EndoFb in different regions of the heart by three tracing strategies. **I**, A cartoon image showing the linage tracing results by three strategies. RV, right ventricle; LV, left ventricle; VS, ventricular septum. The quantification data is represented as the mean ± SD; \**P* < 0.05; NS, non-significant; n = 5 mice. Scale bars: yellow, 1 mm; white, 100 µm.

**Figure 3.**
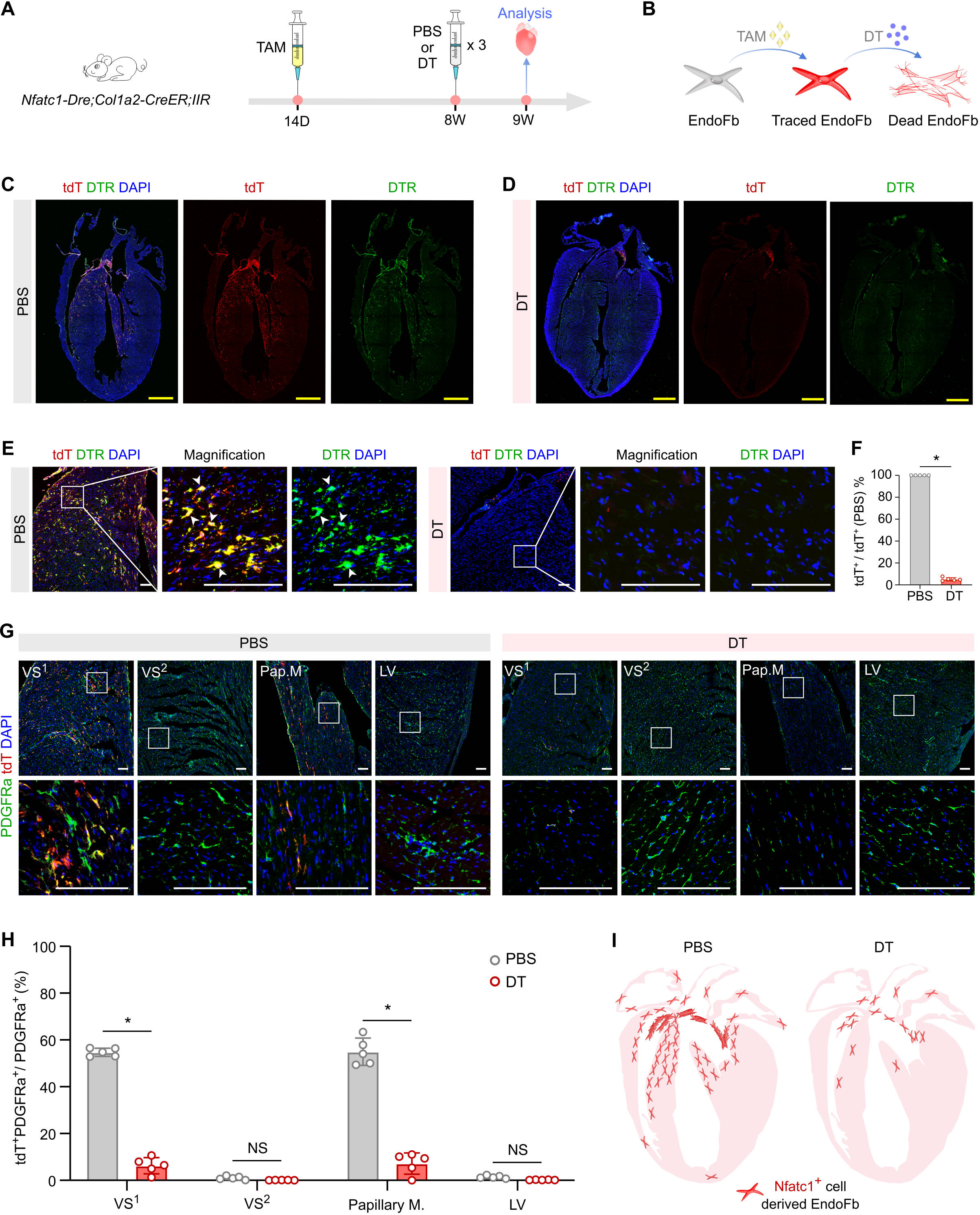
The IIR system enables effective cell ablation. A, A schematic showing the experimental design. **B,** A cartoon showing that the *IIR* enables genetic tracing and ablation of EndoFbs. **C,D,** Immunostaining for DTR and tdT on adult heart sections collected from PBS- (**C**) or DT-treated (**D**) mice. **E,** Immunostaining for DTR and tdT on adult heart sections collected from PBS- or DT-treated mice. **F,** Quantification of the percentage of the tdT^+^ cells in hearts collected from mice treated with PBS or DT. **G,** Immunostaining for tdT and PDGFRa on the different parts of the heart collected from mice with PBS- or DT-treatment. **H,** Quantification of the percentage of PDGFRa^+^ fibroblasts expressing tdT in different regions of hearts collected from mice treated with PBS or DT. **I,** Cartoon images showing the distribution of EndoFbs in hearts collected from mice treated with PBS or DT. LV, left ventricle; VS, ventricular septum; Papillary M., papillary muscle. The quantification data is represented as the mean ± SD; \**P* < 0.05; NS, non-significant; n = 5 mice. Scale bars: yellow, 1 mm; white, 100 µm.

**Figure 4.**
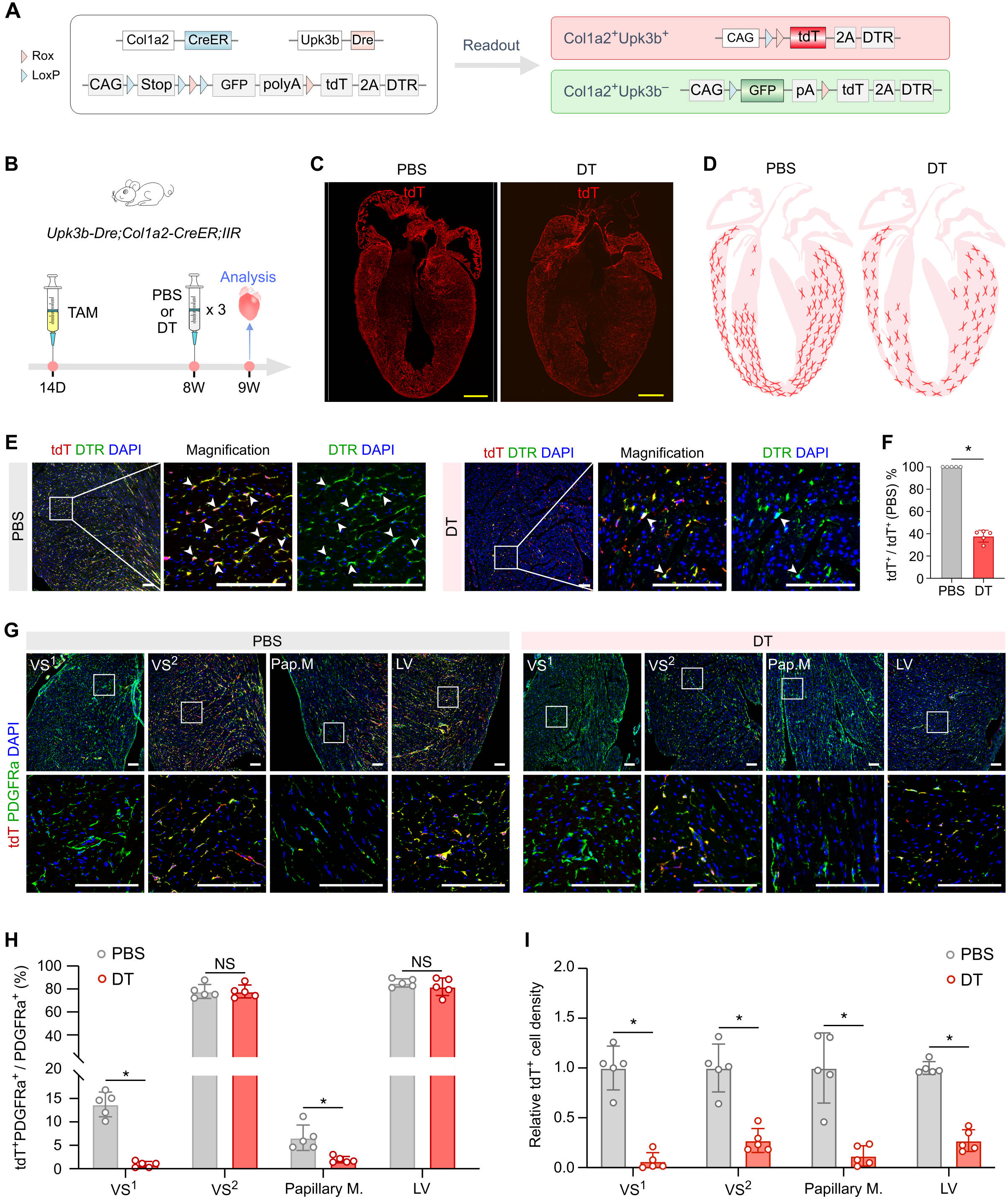
Genetic tracing and ablation of EpiFbs by *IIR*. A, A schematic showing the mouse- crossing strategy. **B,** A schematic showing fluorescent reporter readout of EpiFbs and other fibroblast populations by using *Col1a2-CreER;Upk3b-Dre;IIR* mice. **C,** Immunostaining for tdT on adult heart sections collected from PBS- or DT-treated mice. **D,** Cartoon images showing the distribution of EpiFbs in hearts collected from mice treated with PBS or DT. **E,** Immunostaining for DTR and tdT on adult heart sections collected from PBS- or DT-treated mice. **F,** Quantification of the percentage of the tdT^+^ cells in DT-hearts to that of PBS-hearts. **G,** Immunostaining for tdT and PDGFRa on the different parts of the heart sections collected from mice with PBS- or DT-treatment. **H,** Quantification of the PDGFRa^+^ fibroblasts expressing tdT in different regions of hearts collected from mice treated with PBS or DT. **I,** Quantification of the relative tdT^+^ cell density per area in different parts of hearts collected from mice treated with PBS or DT. LV, left ventricle; VS, ventricular septum; Papillary M., papillary muscle. The quantification data is represented as the mean ± SD; \**P* < 0.05; NS, non-significant; n = 5 mice. Scale bars: yellow, 1 mm; white, 100 µm.

## RESULTS

### Identification of *Nfatc1-Dre*-mediated Ectopic Lineage Tracing

To reassess if Nfatc1 is specifically expressed by EndoFbs within the heterogeneous fibroblast populations of the adult heart, we first used the previously described *Nfatc1-Dre;Col1a2- CreER;R26-RL-GFP* mice^28^ to trace EndoFb (Figure 1A). We collected hearts at 2 weeks after the last injection of Tam and performed immunostaining for GFP and fibroblast marker PDGFRa on frozen heart sections (Figure 1B,C). We found GFP^+^ cells enriched in the upper part of the ventricular septum (VS) and papillary muscle of the left ventricle (LV) (Figure 1C,D), and observed some sporadically distributed GFP^+^ cells in other regions of the heart (Figure 1C,D), consistent with the previous study showing a subset of GFP^+^PDGFRa^+^ fibroblasts in some sub- epicardial regions.^28^ However, we could not definitively determine if these GFP^+^ cells were EndoFbs that migrate into different parts of the heart or if this represented an issue of ectopic labeling by *Nfatc1-Dre* in other fibroblast populations rather than EndoFbs. One method to solve such a problem is to use an alternative promoter specific for endocardial cells that is not activated at any time in all adult cardiac fibroblasts.

Considering that Cdh5 labels endocardial endothelial cells, and adult endothelial cells do not contribute to fibroblasts or vice versa,^30, 43^ we attempted to examine whether the pattern of EndoFbs labeled by *Cdh5-Dre*^41^ would be similar as that traced by *Nfatc1-Dre*. We, therefore, generated the *Cdh5-Dre;Col1a2-CreER;R26-RL-GFP* mouse for comparison with the *Nfatc1- Dre;Col1a2-CreER;R26-RL-GFP* mouse (Figure 1E). We adopted the same experimental strategy for Tam induction and tissue analysis (Figure 1F). Immunostaining for PDGFRa and GFP on frozen heart sections revealed highly enriched GFP^+^ fibroblasts in the upper part of the VS and papillary muscles of LV in the *Cdh5-Dre;Col1a2-CreER;R26-RL-GFP* mice (Figure 1G,H), consistent with that of the *Nfatc1-Dre;Col1a2-CreER;R26-RL-GFP* mice (Figure 1C,D). However, we observed a noticeable reduction of GFP^+^ fibroblasts in other heart regions of the *Cdh5-Dre* tracing mice, such as the lower part of the right ventricular (RV) myocardium or the outer part of the LV myocardium (Figure 1I-K), compared with that of the *Nfatc1-Dre* tracing mice. Data quantification showed a significant reduced percentage of GFP^+^ fibroblasts in RV myocardium, the lower part of VS, and the LV myocardium in the *Cdh5-Dre* compared with that of the *Nfatc1-Dre* tracing mice (Figure 1L). The side-by-side comparison clearly demonstrated that *Nfatc1-Dre* labels a subset of non-endocardium derived fibroblasts in the adult heart (Figure 1D).

With reference to the ectopic labeling of non-endocardium derived fibroblasts in the *Nfatc1-Dre* tracing mice, we reassessed the preferential contribution of EndoFbs to cardiac fibrosis in *Cdh5- Dre* tracing mice after pressure overload-induced heart injury as previously reported^28^. We performed transverse aortic constriction (TAC) in adult hearts and collected the hearts at 4 (4d) or 7 days (7d) post TAC (Figure 1M). In *Cdh5-Dre;Col1a2-CreER;R26-RL-GFP;R26-LSL-tdT* mice, EndoFbs are labeled as GFP^+^ and all fibroblasts regardless of their origins are labeled as tdT^+^ (Figure 1M). Whole-mount fluorescence imaging showed increased GFP^+^ signals in heart samples at 4d and 7d after TAC compared with sham operation (Figure 1N), which was accompanied by increased fibrosis as evaluated by Sirius Red staining (Figure 1O). Immunostaining for GFP and tdT on frozen heart sections at 4d and 7d after TAC revealed remarkably increased GFP^+^tdT^+^ fibroblasts in the upper part of VS and papillary muscle of LV (Figure 1P,Q). In line with the previous work, these data using the *Cdh5-Dre* tracing mice also demonstrated that EndoFbs preferentially responded to TAC with significant expansion after injury. The current data also revealed that some *Nfatc1-Dre* labeled fibroblasts in other regions, such as the RV myocardium or the outer part of the LV myocardium were not EndoFbs likely due to ectopic lineage tracing by intersectional genetic approach.

### Design and Generation of *IIR* Strategy

The above example shows the caveat of using a constitutively active recombinase such as Dre with CreER to target a particular cell population derived from a specific developmental origin. By design, we used A promoter to drive Dre (A-Dre) for targeting A^+^ cells, and B promoter to drive CreER (B-CreER) for targeting B^+^ cells after Tam treatment. By using the R26-RL- Reporter, A^+^B^+^ cells would be labeled as Reporter^+^, and A^−^B^+^ as Reporter^−^ (Figure 2A). During a long time window to study cellular response in homeostasis or after injury, transient activation of A-Dre in A^−^B^+^ cells would label them as Reporter^+^, yielding non-specific labeling (Figure 2A). Unless tracing is controlled by a well-characterized, cell lineage-specific marker such as Cdh5, it is more often limited by a lack of cell specific marker and knowledge about its expression duration in a cell type of interest. It could then be difficult to discern if the constitutively active recombinase would ectopically label other cells transiently expressing the “marker” gene during development, leading to data misinterpretation or overestimation in the contribution of a cell type to adult tissue homeostasis, injury, and diseases. To address this issue, we iterated the above R26-RL-Reporter by incorporating an interleaved strategy (loxP-rox-loxP-rox). We attempted to generate a CAG-loxP-Stop-loxP-rox-loxP-Stop-rox-Reporter line, combining both intersectional and interleaved strategies, named as IIR (Figure 2A). In this IIR mouse, A-Dre;B-CreER would label A^+^B^+^ cells as Reporter^+^, similar to the intersectional strategy as aforementioned. Different from the previous strategy, CreER would also remove the first rox site in A^−^B^+^ cells, creating the loxP-Stop-rox-Reporter allele in A^−^B^+^ cells (Figure 2A). Even if A-Dre is activated transiently in A^−^B^+^ cells, Dre could not recombine loxP-Stop-rox thereafter, preventing ectopic labeling of A^−^ B^+^ cells. As a result, we could still use a constitutively active Dre in combination with an inducible CreER for specific labeling of A^+^B^+^ cells without ectopic labeling of A^−^B^+^ cells, which mitigates the concerns of transient activation of Dre during development or disease progression.

We next generated an IIR mouse line by knocking CAG-loxP-Stop-loxP-rox-loxP-GFP-rox-tdT- P2A-DTR into the Rosa26 locus through homologous recombination using CRISPR/Cas9 as previously described.^44^ We then crossed *Nfatc1-Dre;Col1a2-CreER* with *IIR* to generate *Col1a2- CreER;Nfatc1-Dre;IIR* triple knock-in mice, expecting that EndoFbs would be specifically labeled as tdT (Figure 2B). We treated *Nfatc1-Dre;Col1a2-CreER;R26-RL-GFP* (as a control for comparison) and *Nfatc1-Dre;Col1a2-CreER;IIR* mice with Tam at 14 days after birth, and collected hearts for analysis at 8 weeks old (Figure 2C). Immunostaining for GFP and tdT on frozen heart sections revealed that EndoFbs were highly enriched in the upper part of VS and papillary muscle of LV in both mice (Figure 2D-G). On the other hand, the number of labeled EndoFbs in other parts of the heart was significantly reduced in mice containing IIR compared with that containing *R26-RL-GFP* reporter (Figure 2F-H), confirming that the IIR strategy can significantly reduce ectopic labeling of fibroblasts by *Nftac1-Dre*. Additionally, we also compared the EndoFbs labeled by the IIR strategy with that in *Cdh5-Dre;Col1a2-CreER;R26- RL-GFP* mice, and found no significant difference in the proportion of EndoFb labeling in different parts of the hearts of these two mice (Figure 2H). Our findings demonstrated that IIR strategy effectively prevents ectopic labeling by *Nftac1-Dre* (Figure 2I). Taken together, we presented an effective strategy to mitigate ectopic lineage tracing driven by a constitutively active Dre mouse line.

### Genetic Ablation of EndoFbs by *IIR*

This new *IIR* mouse line contains DTR in tandem with tdT to ensure the recombined cells to express tdT and DTR simultaneously, which enables genetic ablation of tdT^+^ cells by DT treatment. We treated *Nftac1-Dre;Col1a2-CreER;IIR* mice with Tam at 14 days old, administered DT or PBS at 8 weeks old, and collected hearts for analysis at 9 weeks old (Figure 3A). We expected that tdT^+^ EndoFbs would be depleted by DT, leading to reduced tdT^+^ cells in the DT-treated than PBS-treated heart (Figure 3B). Immunostaining for tdT and DTR on *Nftac1- Dre;Col1a2-CreER;IIR* heart sections showed co-localized tdT^+^ and DTR^+^ signal patterns in PBS-treated hearts (Figure 3C), but rare tdT^+^ or DTR^+^ signals were detected in DT-treated hearts (Figure 3D). Magnified images of the upper part of VS clearly showed a significant reduction of tdT^+^DTR^+^ EndoFbs in DT-treated hearts compared with that of PBS-treated ones (Figure 3E,F), suggesting efficient genetic ablation of EndoFbs in *IIR* mice after DT treatment. In addition, tdT^+^ EndoFbs were also remarkably reduced in other parts of the heart in DT-treated mice compared with the PBS-treated group (Figure 3G,H). Taken together, our findings suggested that the IIR strategy enables efficient genetic ablation of labeled cells (Figure 3I), permitting functional evaluation of cells of interest in the pathophysiological process.

### *IIR* Enables Genetic Labeling and Ablation of EpiFbs

In the mouse heart, the majority of fibroblasts are derived from epicardium through epithelial-to- mesenchymal transition during heart development.^45^ Epicardial cells delaminate from the heart surface and migrate into the underlying myocardium, where they differentiate into fibroblasts, smooth muscle cells, pericytes, adipocytes, and a subset of coronary vascular endothelial cells.^31, 32, 46, 47^. To specifically label epicardium-derived fibroblasts (EpiFbs), we used epicardium- specific Dre and *Col1a2-CreER* mice and crossed them with *IIR* instead of *R26-RL-GFP* reporter (Figure 4A). We first used the gene promoter of uroplakin 3B (Upk3b) to drive Dre recombinase and generated *Upk3b-Dre* knock-in mouse line to label epicardial cells and all their descendants. In *Upk3b-Dre;Col1a2-CreER;IIR* mice, epicardium-specific Dre removes the rox-flanked sequence and fibroblast-specific CreER removes the loxP-flanked sequence after Tam, leading to tdT and DTR expression in EpiFbs (Figure 4A). The advantage of using *IIR* in this study is that Tam-induced Cre-loxP recombination removes one rox site, resulting in the generation of loxP-GFP-rox allele in non-epicardium derived fibroblasts (Figure 4A). Such allele would prevent any further Dre-rox recombination, even if there is unexpected *Upk3b-Dre* activation in non-target cells during the long time window of adult homeostasis or after cardiac injury, ensuring no ectopic lineage tracing from the constitutively active Dre recombinase.

To examine if this IIR system could be used for tracing EpiFbs as designed, we treated *Upk3b- Dre;Col1a2-CreER;IIR* mice with Tam at 14 days old and injected PBS or DT at 8 weeks old, and collected hearts for analysis at 9 weeks old (Figure 4B). Immunostaining for tdT on frozen heart sections of PBS-treated mice showed that tdT^+^ EpiFbs were densely distributed in most parts of the heart except for the upper part of VS and papillary muscle of LV (Figure 4C, D). In fact, this unique distribution pattern was complementary to that of EndoFbs. DT injection resulted in a significant reduction of tdT^+^ EpiFbs throughout the heart (Figure 4C, D). Immunostaining for tdT and DTR on frozen heart sections collected from *Upk3b-Dre;Col1a2- CreER;IIR* mice showed a considerable reduction of tdT^+^DTR^+^ EpiFbs in DT-treated hearts compared with that of PBS-treated ones (Figure 4E,F). While the total number of tdT^+^ EpiFbs was reduced in different regions of the hearts, we next asked if the percentage of the remaining PDGFRa^+^ cells expressing tdT would be changed in these regions. By examining different regions of the heart, we found that the percentage of PDGFRa^+^tdT^+^ fibroblasts among total fibroblasts was significantly reduced in the upper part of VS (VS^1^) and papillary muscles of LV, but remained similar to that of the PBS-treated hearts in the lower part of VS (VS^2^) and myocardial wall of LV (Figure 4G,H). The observation reflects the potential compensation by other fibroblast populations (most likely EndoFbs) to replace the loss of EpiFbs in specific regions, such as VS^1^ and papillary muscle of LV. In VS^2^ and myocardial wall of LV where EpiFbs were predominantly located, the percentage of PDGFRa^+^tdT^+^ fibroblasts among total fibroblasts was not significantly reduced, indicating that the small number of EndoFbs were less likely to compete with the remaining EpiFbs for expansion in these regions (Figure 4H). Of note, the density of EpiFbs in all different regions of the heart was reduced significantly after DT treatment, compared with that of the PBS-treated group (Figure 4I), indicating the high efficiency of genetic ablation in EpiFbs by IIR system.

## DISCUSSION

Here, we developed an interleaved intersectional reporter (IIR) strategy to resolve the ectopic labeling issue associated with unwanted activation of dual recombinases Dre and CreER that are broadly used in genetic lineage tracing studies. The application of dual recombinases-mediated genetic lineage tracing has improved the precision of cell fate mapping study. However, some technological caveats remain to be addressed when a constitutively active recombinase such as Dre is used in combination with an inducible homologous recombinase such as CreER. The unexpected despite transient activation of Dre in some “unwanted” cells may lead to Dre-rox recombination, resulting in ectopic cell labeling and, therefore, data misinterpretation in lineage tracing studies. We designed an IIR strategy to use CreER to effectively prevent such Dre-rox recombination by removing one rox site by CreER in the early stage, such as the neonatal stage, before analysis in the adult tissues. In addition, “Dre + CreER” combination allows for broad utilization of both constitutive and inducible homologous recombinases in cell fate study. To showcase the capability and effectiveness of IIR strategy, we distinctively labelled EndoFbs and EpiFbs in the mouse heart, respectively. Moreover, we can couple lineage tracing with genetic cell ablation for cell fate study and functional evaluation simultaneously *in vivo*.

In the mammalian heart, fibroblasts are heterogeneous^48, 49^ and are mainly derived from two major developmental origins: epicardium and endocardium.^34^ While previous studies have reported that developmental heterogeneity of cardiac fibroblasts does not predict their pathological proliferation and activation,^33^ more specific lineage tracing and genetic manipulation experiments demonstrated that EndoFbs represent a unique fibroblast population that preferentially proliferate and expand in response to pressure overload,^28^ highlighting the importance of using genetic technology with enhanced precision to study the contribution of subpopulations of cells to cardiovascular disease. To label EndoFbs, *Nfatc1-Dre*, proposed as endocardium-specific, was used in combination with *Col1a2-CreER* and *R26-RL-GFP* reporter. While GFP^+^ fibroblasts were highly enriched in the upper part of VS and the papillary muscle of LV, we also observed a subset of PDGFRa^+^ fibroblasts expressing GFP in sub-epicardial regions that are far away from the papillary muscle of the LV.^28^ We interpreted such an unexpected labeling as resulted from transient activation of *Nfatc1-Dre* in other fibroblast subsets after birth. However, it is also possible that a subset of EndoFbs may migrate away to reach the sub-epicardial region where they reside and expand in the adult stage. We could not distinguish these two possibilities with *Nfatc1-Dre* itself, as it is technically challenging to prevent transient activation of Nfatc1 during development by targeting its promoter driving Dre recombinase in the adult heart. We employed a positive control by designing endothelial cell-specific promoter Cdh5 to drive Dre, that labels endocardial endothelial cells in addition to coronary vascular endothelial cells. Given that adult endothelial cells do not contribute to fibroblasts and vice versa,^30, 43^ we could use endothelial cell-specific driver Cdh5-Dre to label EndoFbs. In *Cdh5- Dre*;*Col1a2-CreER;R26-RL-GFP* mouse line, similar distribution patterns of GFP^+^ fibroblasts were observed in the upper part of VS and the papillary muscle of LV, but significantly fewer GFP^+^ fibroblasts were found in other regions compared with the *Nfatc1-Dre* based tracing mice. This comparison confirmed the previous hypothesis that labeling of a subset of GFP^+^ fibroblasts in other regions of the heart is due to non-specific lineage tracing by *Nfatc1-Dre*. For EndoFbs, we found a good control based on *Cdh5-Dre* tracing that only labels EndoFbs but not fibroblasts derived from other origins. However, in many conditions, we may not have positive controls or alternative genetic tools to confirm ectopic labeling in lineage tracing studies. The IIR strategy presented here prevented Dre-rox recombination after target cell labeling at later developmental stage, showing a similar distribution pattern of labeled EndoFbs comparable to that observed in *Cdh5-Dre* tracing mice. This example demonstrates the power of IIR for mitigating non-specific lineage tracing using Dre, which can be broadly used in cardiovascular research to address the contribution and function of a subpopulation of cells in pathophysiological processes.

In this study, we also applied the IIR strategy to specifically label and ablate EpiFb. It is known that epicardium is the major source of cardiac fibroblasts, yet their clear distribution and function remain incompletely understood. Using *Upk3b-Dre*;*Col1a2-CreER;IIR* mice, we found that EpiFbs were distributed in regions complementary to that of EndoFbs. While there is a possibility of ectopic activation of *Upk3b-Dre* in fibroblast populations other than EpiFbs, our IIR could alleviate such concerns as there would be no Dre-rox recombination after Tam injection when *Col1a2-CreER* removes one rox site from IIR allele. Therefore, the genetic labeling of EpiFbs should be specific in the adult heart, offering a precious opportunity to study EpiFbs in cardiac injuries and diseases in the future. As IIR contains both tdT and DTR, we can ablate EpiFbs and quantify the degree of tdT^+^ cell loss simultaneously. Indeed, we depleted the majority of EpiFbs throughout different heart regions and observed a significant reduction of tdT^+^ fibroblasts after DT treatment. While the absolute number of tdT^+^ fibroblasts was reduced, the percentage of tdT^+^ fibroblasts remained relatively stable in most regions of the heart where EpiFbs were the predominant fibroblast residents. Interestingly, in regions where EndoFbs were predominant, removal of EpiFbs led to a significant reduction in the percentage of tdT^+^ fibroblasts among total fibroblasts, suggesting that EndoFbs could have replaced the loss of EpiFbs at homeostasis. In regions where EpiFbs were preferentially located such as the myocardial wall of LV, the few EndoFbs of these regions may not sufficiently replace the loss of EpiFbs. Taken together, our findings suggest competition of heterogenous fibroblast subpopulations derived from different developmental origins may occur in distinct regions of the heart during homeostasis and after injuries, which merits further investigation in the future.

## Acknowledgments

We thank the Shanghai Model Organisms Center, Inc. (SMOC) for mouse generation; and institutional animal facilities for mouse husbandry.

## Author Contributions

M. Han and Z Liu conceived and designed the project. X. Huang performed injury models and bred mice. L. Liu bread mice, performed mouse experiments, and analyzed the data; K. O. Lui, Q. Shu, and B. Zhou interpreted the data, drafted and revised the manuscript. B. Zhou supervised the research.

